# How is Emotional Evidence from Multiple Sources Used in Perceptual Decision Making?

**DOI:** 10.1101/2023.02.28.530147

**Authors:** Hilary H.T. Ngai, Janet H. Hsiao, Christian Luhmann, Aprajita Mohanty, Jingwen Jin

## Abstract

Judging the emotional nature of a scene requires us to deliberately integrate pieces of evidence with varying intensity of emotion. Our existing knowledge about emotion-related perceptual decision making is largely based on paradigms using single stimulus and, when involving multiple stimuli, rapid decisions. Consequently, it remains unclear how we sample and integrate multiple pieces of emotional evidence deliberately to form an overall judgment. Findings from non-emotion rapid decision-making studies show humans down-sample and downweight extreme evidence. However, deliberate decision making may rely on a different attention mode than in rapid decision making; and extreme emotional stimuli are inherently salient. Given these critical differences, it is imperative to directly examine the deliberate decision-making process about multiple emotional stimuli. In the current study, human participants (N=33) viewed arrays of faces with expressions ranging from extremely fearful to extremely happy freely with their eye movement tracked. They then decided whether the faces were more fearful or happier on average. In contrast to conclusions drawn from non-emotion and rapid decision-making studies, eye movement measures revealed that participants attentionally sampled extreme emotional evidence more than less extreme evidence. Computational modeling results showed that even though participants exhibited biased attention distribution, they weighted various emotional evidence equally. These findings provide novel insights into how people sample and integrate multiple pieces of emotional evidence, contribute to a more comprehensive understanding of emotion-related decision making, and shed light on the mechanisms of pathological affective decisions.

## Introduction

In daily life, an emotional event often consists of a plethora of emotional stimuli. When forming an overall judgment regarding the emotion in such contexts observers will integrate multiple pieces of evidence with varying intensity of emotion, a process that takes some time. However, emotion perception research has primarily relied on the identification of an isolated emotional stimulus presented only briefly (Calvo & Nummenmaa, 2016; Wilhelm et al., 2014). Consequentially, the process by which people sample and integrate varied pieces of emotional evidence purposefully and deliberately to form the final perceptual decision remains unclear. Understanding the mechanisms of such multi-evidence decision making is critical for adaptive interactions with our complex environments because errors in this process can yield biased impressions with detrimental personal and social consequences. For example, over-sampling and overweighting highly emotional stimuli may bias one to make extreme choices whereas down-sampling and downweighting them may lead to decisions that put one’s well-being at risk.

Perceptual decision making summarizing multiple stimuli is constrained by our perceptual system (Broadbent, 1953; Luck & Vogel, 1997). It is theorized that observers overcome visual capacity constraints by using two attentional strategies (Baek & Chong, 2020). Humans can extract the summary statistics across multiple pieces of evidence presented only briefly, forming an ensemble representation (Alvarez, 2011; Ariely, 2001; Chong & Treisman, 2003; Watamaniuk et al., 1989; Whitney & Yamanashi Leib, 2018). Such ensemble perception relies on distributed attention spanning all stimuli (Alvarez & Oliva, 2008; Elias et al., 2017; Li et al., 2016; Wolfe et al., 2015). In contrast, the focused attention mode, used for recognizing a specific visual object, deals with the visual capacity constraints by selecting and processing salient or relevant stimuli and filtering out irrelevant stimuli (Carrasco, 2011; Chun, Golomb, & Turk-Browne, 2011). Some researchers have argued that observers use both distributed and focused attention modes when dealing with a limited perceptual capacity depending on the goal (Baek & Chong, 2020).

The key limitation of perceptual averaging research is that it has relied mainly on stimuli that are not inherently motivationally salient and decision-making tasks that are rapid (e.g., stimuli are available for less than a fraction of a second). This research has shown that when arriving at an overall decision regarding multiple stimuli, observers mitigate the influence of extreme evidence on perceptual judgments, like a statistician excludes outlying data points. For example, in rapid decision making, people engage in “robust averaging”, downweighting outlying or extreme evidence because it is considered unreliable or untrustworthy (de Gardelle & Summerfield, 2011; Epstein et al., 2020; Haberman & Whitney, 2010; Huber, 2004; Li et al., 2017; Vandormael et al., 2017; Whitney & Yamanashi Leib, 2018). Furthermore, inlying (closer to average) compared to outlying evidence attracts more visual attention (Vandormael et al., 2017), indicating robust sampling in addition to robust weighting. Together, these findings suggest that ensemble perception is a rapid, automatic, and implicit process that is relatively immune to outliers (Haberman et al., 2009; Haberman & Whitney, 2007, 2010). Research examining decisions regarding multiple emotional stimuli has been sparse, with existing knowledge mainly from rapid perceptual tasks (Goldenberg et al., 2020; Haberman & Whitney, 2010; Sun & Chong, 2020; Wolfe et al., 2015).

The factors involved in making deliberate decisions – without the duress of time – regarding multiple emotional stimuli are likely to be different than those involved in rapid extraction of summary information regarding non-emotional stimuli (Bogacz et al., 2010; Kiani et al., 2008; Pessoa, 2009; Phelps et al., 2014). Specifically, with the time constraint lifted, observers can carry out *purposeful* and *active* sampling and *resampling* of the evidence space by shifting focused attention to properly register each stimulus (Gidlöf et al., 2013; Itti & Koch, 2000; Kietzmann et al., 2011; Posner, 1980; Rahn et al., 2016; Wyart & Tallon-Baudry, 2008). In this context, visual attention allocation can be influenced by the saliency and value of the stimuli as well as the goal of the observer, which can be suitably measured using eye movement data (Christie et al., 2018; Grebitus et al., 2015; Krajbich et al., 2010; Krajbich & Rangel, 2011; Kulke, 2019; Sepulveda et al., 2020). Due to evolutionary saliency, emotional stimuli tend to capture attention in a “bottom-up”, stimulus-driven manner (Feldman, 1995; Orquin & Loose, 2013; Posner et al., 2005). Indeed, in free-viewing visual search tasks, eye movements tend to be attracted towards locations where emotionally salient stimuli are present (Calvo & Nummenmaa, 2016; Kulke, 2019; Mogg & Bradley, 1999; Nummenmaa et al., 2006; Towal et al., 2013; Võ & Henderson, 2010). Thus, it is likely that outlying (i.e., extreme) compared to inlying (i.e., closer to the mean) emotional stimuli would attract more visual attention. When considering evidence weighting, there are findings from two alternative tasks showing that visual attention is associated with decision preference, such that decision is biased towards choosing the option that attracted more eye gaze (Armel et al., 2008; Hertwig & Erev, 2009; Kaanders et al., 2022; Pärnamets et al., 2016; Thomas et al., 2019). Visual attention favoring the inlying than outlying evidence is also linked to the weights in non-emotion decision-making (Vandormael et al., 2017). However, emotion-related decision making based on perceptual averaging of multiple stimuli has not been examined using such free-viewing paradigms; thus, it remains to be tested whether visual attention translates into decision weight in this context.

We used a free-viewing perceptual decision-making task, eye tracking, and computational modeling to elucidate how people sample and integrate multiple pieces of emotional evidence. Eye movement data were recorded as participants viewed multiple emotional faces ranging from extremely fearful to extremely happy before deciding whether the array of faces was more fearful or happier on average. We hypothesized that faces showing more extreme emotions would attract more attention. This would manifest as a U-shape pattern of eye movements as a function of faces ordered from extremely fearful to extremely happy. Moreover, using computational modeling we tested the hypothesis that this unbalanced attention allocation would translate into a biased weighting such that stimuli attracted more attention would be weighted higher in forming the perceptual decision.

## Methods

### Participants

Forty-six participants (*Mean Age* = 20.4 years old, *SD* = 3.0 years old; Women = 71.7%, Chinese & Other Asian = 95.7%) from the University of Hong Kong were recruited through posters and from the Psychology Department Research Participant Pool. Given that there was no existing research using the same task paradigm, the sample size was determined based on an influential study investigating non-emotion-related rapid decision-making of multiple stimuli(de Gardelle & Summerfield, 2011). In this study, the sample sizes for four experiments were 30, 14, 16 and 24 respectively. We took a conservative approach and decided on the target sample size to be 30.

Data from 13 participants were discarded due to lack of completion (N = 4), technical failure (N = 5) and being outliers in eye movement measurements (N = 4, see Eye Movement Data Processing). Thus, the final dataset consisted of 33 participants. All participants provided written informed consent, reported fluency in English, and had normal or corrected-to-normal vision. The study was approved by the Human Research Ethics Committee of the University of Hong Kong. Participants were either given partial course credit or were compensated 50HKD per hour.

## Stimuli

Image stimuli consisted of 10 (5 fearful and 5 happy) faces selected from 5 Asian actors (2 female) from the HKU Face Database (Zhang et al., 2019). Asian face actors were chosen to match the local sociocultural context (Masuda et al., 2008). Fearful and happy faces were chosen because these emotions are both high arousal yet of opposite valence (Watson et al., 1999). Although both anger and fear are of negative valence, angry faces were not chosen, because anger provokes behavioral changes in the target of that anger (e.g., the participants), eliciting a more complex response (Pichon et al., 2009). The images were first converted from color to grayscale and cropped around the chin and ears (300 x 300 pixels). FantaMorph (Abrosoft Fantamorph Version 5.6.2; www.fantamorph.com) was then used to create an array of faces ranging from fearful to happy. For each face actor, facial features including the corner of eyebrows, shape of the eye, nose, and mouth were matched for their fearful and happy expressions. Then, 99 synthetic images were created by linearly interpolating between the original fearful and happy images, yielding 101 images, where each image depicted varying degrees of fearful/happiness. Morphed images were manually checked and retouched using Adobe Photoshop to ensure that these images were natural-looking (Version 22.1.1). We numerically coded the 101 images, with a morphing unit of 0 corresponding to the face actor’s original fearful face and 100 corresponding to the original happy face. Lastly, the 505 (101 fearful-to-happy images x 5 actors) stimuli were equalized for contrast and luminance using the default functions of the SHINE Toolbox in MATLAB (Willenbockel et al., 2010) (specMatch to equate the Fourier spectra and histMatch to match the luminance histograms).

## Experimental Procedure

All participants were asked to complete two tasks (**Figure 1A-B**), the first being a perceptual decision-making task and the second being a subjective emotion rating task. Both tasks comprised of 75 trials, which were pseudorandomized such that the same face actor was not presented more than four trials in a row to avoid local learning effects. On each trial, eight faces belonging to the same actor were randomly assigned into eight placeholders in an annulus format while a ninth face was presented in the center of the annulus. The eight images in the annulus formed the multi-evidence face set, with varying emotion intensities (i.e., morphing units) while the image in the center depicted 50% fearfulness and 50% happiness for that face actor based on the morphing unit. Here onwards, we use objective emotion value (OEV) to refer to the morphing unit. OEV of the eight faces were determined for each trial using a Gaussian distribution with a mean of 50 (i.e., the midpoint of the fearful to happy spectrum). To create trials with variable coverage of OEV, three levels of variance (small, medium, and large) were used in the Gaussian distribution, with 25 trials of each level of variance.

**Figure 1.**
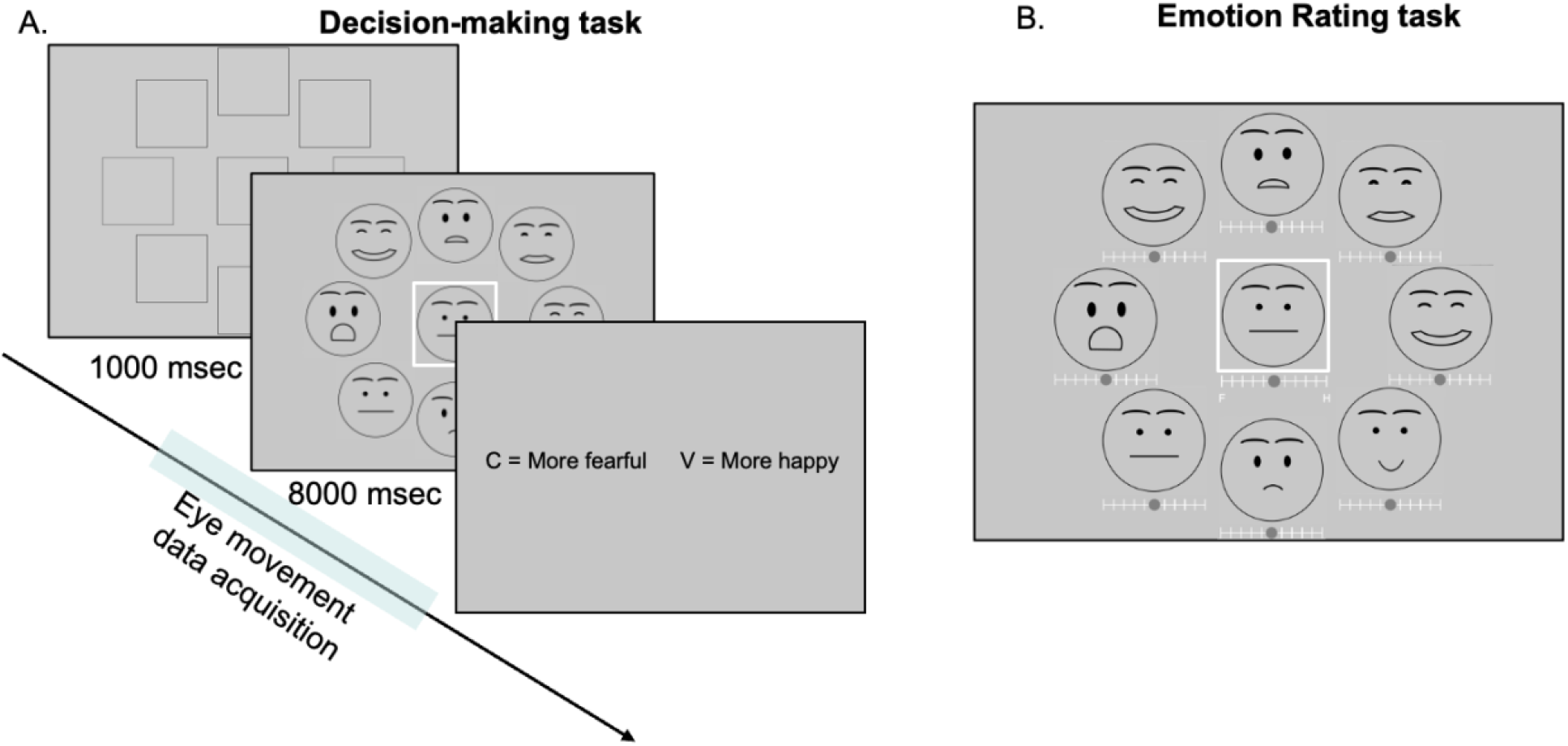
Experiment tasks. (**A**) Trial timeline of the decision-making task. After empty placeholders, eight faces were shown for 8000-msec, during which eye movements were recorded. A ninth reference face was placed in the center. Participants then made a binary decision on whether the eight faces were more fearful or happier on average than the reference face. **(B)** Sample trial for the subjective emotion rating task. Instead of making a binary decision, participants rated each individual face on a scale of “Fearful to Happy” without time constraints. In both **A** and **B,** the center face was that trial’s particular face actor/actress’s face with the OEV of 50. Participants were instructed to reference that face when making a decision and rating.

### The decision-making task

In the decision-making task **(Figure 1A)**, each trial began with a drift correction check where a solid circle appeared in the center of the screen, and the stimuli were presented after the participants’ fixation was within 1 degree of visual angle. Each trial began with nine empty squares (placeholders) presented for 1000-msec. Then, the set of faces described above replaced their respective placeholders for 8000-msec. Participants were instructed to determine whether the eight faces placed on an annulus format were on an average more fearful or happier than the central face image. Hence, the eight annulus faces provided the evidence space to be integrated for the subsequent decision and the central face simply provided a reference which they could use to make their decision regarding the average emotion. Upon the offset of the face stimuli presentation, participants indicated their overall decision by pressing one of two buttons. Participants could view the evidence space freely during the 8000-msec of stimuli presentation, allowing for purposeful sampling of different pieces of emotional evidence. Their eye movements were tracked during this 8000-msec free viewing. This task was delivered using Experiment Builder (Version 2.2.61) compatible with eye-tracker software.

### The subjective emotion rating task

Since the OEV of faces were unlikely to reflect the subjectively perceived emotion values that may vary across trials and across individuals, we obtained each participant’s own emotion values of the faces using a subjective emotion rating task **(Figure 1B)**. The face image stimuli and trial sequence were the same as used in the decision-making task. But instead of making a binary decision regarding the overall emotion across faces, participants rated each of the eight faces on a scale ranging from 0 (“very fearful”) to 10 (“very happy”) with no time constraint for each trial. Once again, the participants were instructed that the central face image served as the reference of 50% fearful and 50% happy. Ratings obtained from this task were referred to as the subjective emotion value (SEV). Measuring the SEV of each face stimulus within the face array context in each trial allowed us to isolate the role of SEV from the role of evidence sampling and weighting. This task was delivered using PsychoPy (Version 3.2.4).

## Apparatus

Eye movement of each participant’s dominant eye was recorded using an EyeLink 1000 eye-tracker (SR Research Ltd.). Participants sat 60-cm in front of a 22ʺ monitor (1024 × 768 pixels). Using pupil and corneal reflection, the system has a sampling rate of 1000 Hz which was suitable for monocular sampling. The default settings of Eyelink (i.e. saccade motion threshold of 0.1° visual angle, saccade acceleration threshold of 8000°/s^2^ and saccade velocity threshold of 30°) were used during data collection. At the beginning of the perceptual decision-making task, the default nine-point calibration and validation were conducted and repeated until the drift correction error was less than 1 degree of visual angle. All participants used a chin rest to reduce head movements.

## Eye Movement Data Processing

When analyzing the eye movement data, we considered both the spatial (i.e., fixation location) and temporal dimensions (i.e., duration, and the sequence of transition amongst the fixation locations). Fixation count and gaze duration were calculated for each region of interest (ROI), which corresponded to the nine placeholders within each trial. The nine placeholders were defined by their corresponding coordinate vertices, covering a 180-by-180-pixel area. The ROIs for transition probability were optimized for the Eye Movement analysis with Hidden Markov Model (EMHMM) estimation (detailed below). Quality check was conducted for eye movement data by finding the mean number of fixations of all participants and then detecting outliers by using the mean ± 2 SD as the threshold. Four participants were discarded as they had more than 20% outlier trials.

### Transition probability estimated with Hidden Markov Models (EMHMM)

Transition probability is a measure to describe the likelihood of a saccade towards an ROI, based on the location of the previous fixation. Transition probability attaches importance towards the trajectory of eye movement in a trial by taking into account both spatial and temporal information of a participant’s visual attentional pattern (Chuk et al., 2014). The inclusion of transition probability thus provided valuable information in addition to fixation count and gaze duration. To estimate the transition probability, we employed a data-driven, machine-learning approach using the EMHMM MATLAB toolbox (version 0.77). EMHMM uses Hidden Markov Models (HMM) to analyze the observable time-series data provided from eye-tracking. EMHMM assumes that the ROI of each fixation is dependent only on the previously viewed ROI. The hidden states of an HMM corresponded to the ROIs, and the observable fixation data were modeled as arising from an underlying dynamic process of the sequence of ROIs viewed (Chan et al., 2018; Chuk, Chan, et al., 2017; Chuk, Crookes, et al., 2017). Since our task included a natural set of nine ROIs of each trial, we predefined the spatial ROIs using the area of each face’s location defined by the coordinates of the center with a radius of 90 pixels (i.e., 4 SD of a 2-dimensional Gaussian distribution) (Cho et al., 2022a; Cho et al., 2022b; Chuk et al., 2019). The prior probabilities across the nine ROIs were initialized as a flat distribution. Then, using the fixation point sequence from the eye movement data, the HMM estimated the probability of starting at a particular ROI (i.e. estimated prior probability) and the probability of fixations transitioning between a pair of ROIs (i.e. transition probability) (Chuk et al., 2014). All analyses regarding the eye movement data were conducted in the environment of MATLAB (Release 2020b; The MathWorks, Natick, MA).

## Statistical Analyses

### Attention allocation using eye movement measurements

To investigate how participants allocated their overt visual attention across the evidence-bearing face-set, the mean proportion of fixation counts, proportion of gaze durations, and marginal transition probabilities to each ROI were analyzed.

For fixation count and gaze duration, we first grouped fixations by ROIs based on coordinate location, excluding fixations that fell outside the nine ROIs. This was done for each trial. Then, for each of the eight evidence-bearing ROIs (excluding the central reference image), the proportion of fixation counts, and gaze duration values were computed within each trial. Finally, following previous research (de Gardelle & Summerfield, 2011), for each trial these fixation counts and gaze durations for the eight ROIs were reordered from the most fearful to the happiest based on the SEVs. Thus, after reordering, ROI1 always referred to the most fearful face of that trial, while ROI8 was the happiest face of that trial. Here onwards, we refer to this factor as the ROI Rank Order. Finally, the mean proportion of fixation counts and gaze duration for each ordered ROI was computed averaging across all trials.

The transition probabilities (reflecting the tendency of shifting the gaze towards a target ROI from any other ROI) to each of the eight evidence-bearing ROIs, were computed by calculating the marginal transition probabilities. For each ROI, this marginal transition probability equaled the sum of all the transition probabilities to the target ROI, weighted by the prior probability of the source ROI **(Equation 1)**. The prior and transition probabilities were the estimated outputs of the EMHMM.

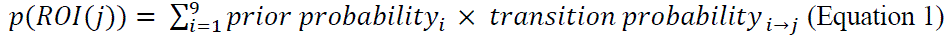

Note that the EMHMM was conducted using the original ROI sequence to preserve fidelity to the natural eye movement sequence. The marginal transition probabilities were then reordered based on the ROI Rank Order outlined above.

For statistical analyses, one-way repeated measures ANOVAs (rmANOVA) with 1 factor (ROI rank order) of eight levels were applied to examine the effect of ROI Rank Order (from the most fearful to the happiest) on each of the three eye movement measures. We were particularly interested to see if they followed the hypothesized U-shaped pattern.

Next, for fixation count and gaze duration, we also conducted analyses examining the first 4000-msec and the second 4000-msec epochs, separately. This was to evaluate whether the visual attention patterns revealed across the 8000-msec was consistently shown in the earlier and latter half of the trial. The same rmANOVA analysis with the ROI Rank Order factor was applied. Given that the marginal transition probability summarizes the trajectory over a trial and splitting the trial could affect the HMM estimation by reducing the number of data points used in the estimation, this epoch-based analysis was not applied to this measure.

### Contribution of attention allocation to evidence weighting

To examine how participants weighted the emotional evidence to form a decision, we compared nine models (see **Table 1**). First, as a baseline (M0), the mean OEV of each trial was used to calculate the decision value (DV), which captured no individual differences as the OEV would be the same for each trial across participants. Then for models 1-2 (M1&2), all pieces of evidence were weighted equally, and the DV was computed using the SEVs of the eight face stimuli in each trial of the emotion rating task. M1 took a categorical approach by dichotomizing each SEV into +1 if it was rated as happier and –1 if it was rated as more fearful before taking the average, with the categorization determined by comparing the SEV to the midpoint. In contrast, M2 took a continuous approach by directly taking the mean value of the eight SEVs, thus M2 retained all the SEV information. The remaining six models tested whether attention allocation – measured using the three eye movement measures – translated into the weights in decision making. Models 3-5 involved dichotomized SEV as in M1. For a given eye movement measure of a given ROI, the eye movement measure was multiplied by –1 if the face was a fearful one and was multiplied by +1 if the face was a happy one. The DV was computed by taking the mean of all the transformed values for the percentage of fixation count (M3), the percentage of gaze duration (M4), and marginal transition probability (M5). Models 6-8 involved continuous SEV as in M2, with DV calculated as a weighted sum of the SEV. Hence, SEV of each face stimulus was directly weighted by one of the eye movement measures: percentage of fixation count (M6), percentage of gaze duration (M7), and the marginal transition probability (M8).

**Table 1:**
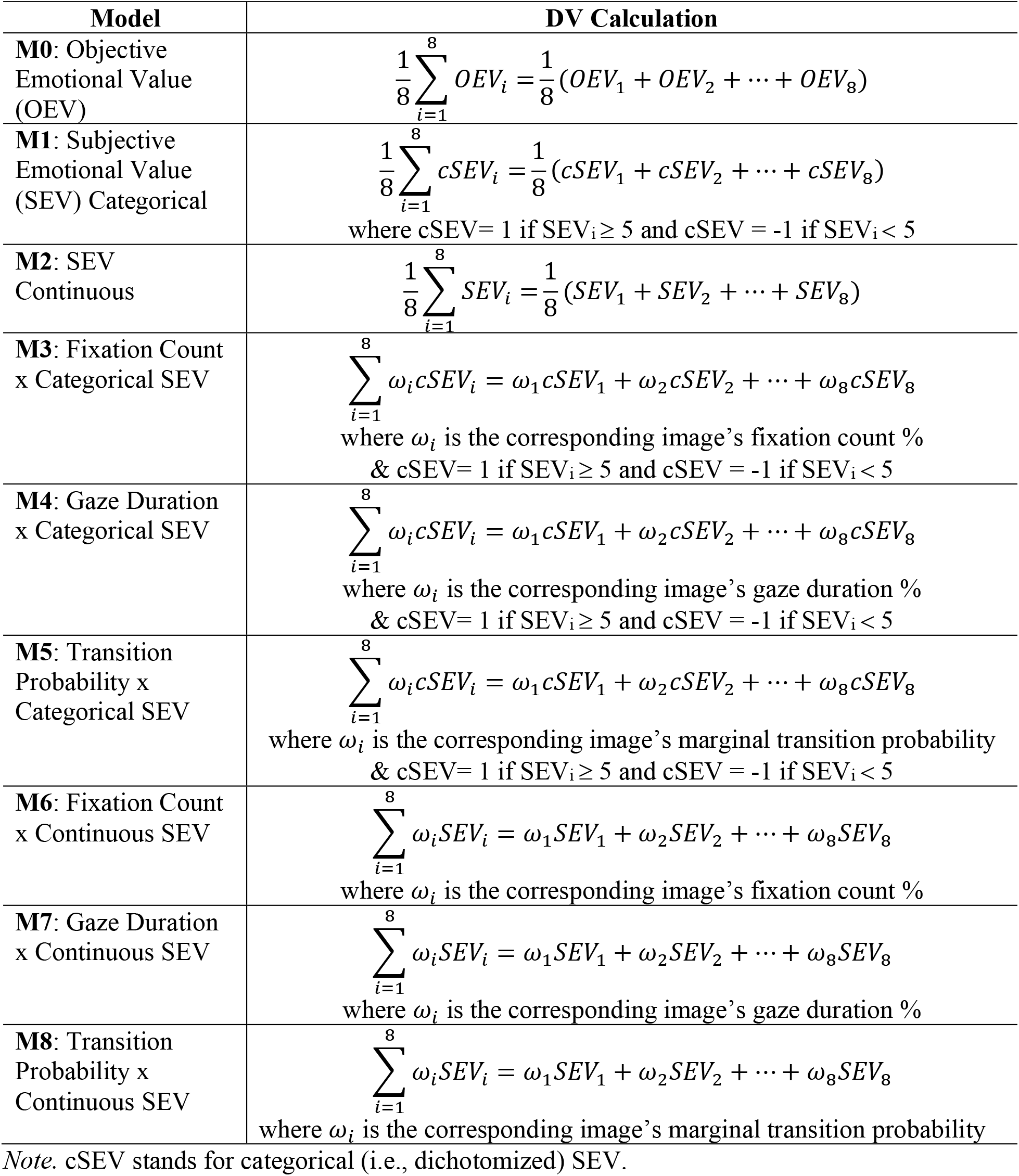
Models and Respective Decision Value (DV) Calculations Per Trial Per Subject.

| **Model** | **DV Calculation** |
| --- | --- |
| **M0:** Objective Emotional Value (OEV) | $\frac{1}{8} \sum_{i=1}^8 OEV_i = \frac{1}{8} (OEV_1 + OEV_2 + \dots + OEV_8)$ |
| **M1:** Subjective Emotional Value (SEV) Categorical | $\frac{1}{8} \sum_{i=1}^8 cSEV_i = \frac{1}{8} (cSEV_1 + cSEV_2 + \dots + cSEV_8)$<br>where $cSEV = 1$ if $SEV_i \geq 5$ and $cSEV = -1$ if $SEV_i < 5$ |
| **M2:** SEV Continuous | $\frac{1}{8} \sum_{i=1}^8 SEV_i = \frac{1}{8} (SEV_1 + SEV_2 + \dots + SEV_8)$ |
| **M3:** Fixation Count x Categorical SEV | $\sum_{i=1}^8 \omega_i cSEV_i = \omega_1 cSEV_1 + \omega_2 cSEV_2 + \dots + \omega_8 cSEV_8$<br>where $\omega_i$ is the corresponding image's fixation count %<br>& $cSEV = 1$ if $SEV_i \geq 5$ and $cSEV = -1$ if $SEV_i < 5$ |
| **M4:** Gaze Duration x Categorical SEV | $\sum_{i=1}^8 \omega_i cSEV_i = \omega_1 cSEV_1 + \omega_2 cSEV_2 + \dots + \omega_8 cSEV_8$<br>where $\omega_i$ is the corresponding image's gaze duration %<br>& $cSEV = 1$ if $SEV_i \geq 5$ and $cSEV = -1$ if $SEV_i < 5$ |
| **M5:** Transition Probability x Categorical SEV | $\sum_{i=1}^8 \omega_i cSEV_i = \omega_1 cSEV_1 + \omega_2 cSEV_2 + \dots + \omega_8 cSEV_8$<br>where $\omega_i$ is the corresponding image's marginal transition probability<br>& $cSEV = 1$ if $SEV_i \geq 5$ and $cSEV = -1$ if $SEV_i < 5$ |
| **M6:** Fixation Count x Continuous SEV | $\sum_{i=1}^8 \omega_i SEV_i = \omega_1 SEV_1 + \omega_2 SEV_2 + \dots + \omega_8 SEV_8$<br>where $\omega_i$ is the corresponding image's fixation count % |
| **M7:** Gaze Duration x Continuous SEV | $\sum_{i=1}^8 \omega_i SEV_i = \omega_1 SEV_1 + \omega_2 SEV_2 + \dots + \omega_8 SEV_8$<br>where $\omega_i$ is the corresponding image's gaze duration % |
| **M8:** Transition Probability x Continuous SEV | $\sum_{i=1}^8 \omega_i SEV_i = \omega_1 SEV_1 + \omega_2 SEV_2 + \dots + \omega_8 SEV_8$<br>where $\omega_i$ is the corresponding image's marginal transition probability |

*Note.* cSEV stands for categorical (i.e., dichotomized) SEV.

Each model assumed that the respective DV generated probabilistic preferences by passing the DV through a conventional softmax function **(Equation 2)**.

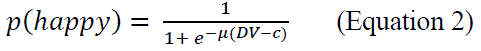

The model contained two free parameters: the value of the DV at which each participant was expected to be indifferent (i.e., *p*(happy)=0.5), *c*, and a determinism parameter that controlled how strongly related choices were to the DV, *μ*. Parameter values were taken as those minimizing the cumulative negative log-likelihood of the choice data. These values were estimated separately for each participant and model using the optimization routines in the SciPy Python package (Virtanen et al., 2020). Following model fitting, the mean of each model’s negative log-likelihood were calculated to summarize which model best described the participants’ task behavior (i.e., by which model yielded the lowest negative log-likelihood). The conclusion of how participants weighted evidence was drawn based on which model explained the choice behavior data the best.

## Preregistration and open science

We pre-registered the experiment on the Open Science Framework (OSF). Pre-registrations, materials are made available on the OSF at https://osf.io/e8jvb. Data can be made available upon request to the corresponding author.

## Results

### Descriptive Results

For the subjective rating task (**Figure 2A**), SEV showed a positive correlation with OEV. The mean correlation across all participants was 0.792 (*SD* = 0.068), which leaves room for SEV to account for unique variance in ratings. This pattern supported the usage of SEV instead of the morphing unit-based OEV in the subsequent data analyses.

**Figure 2.**
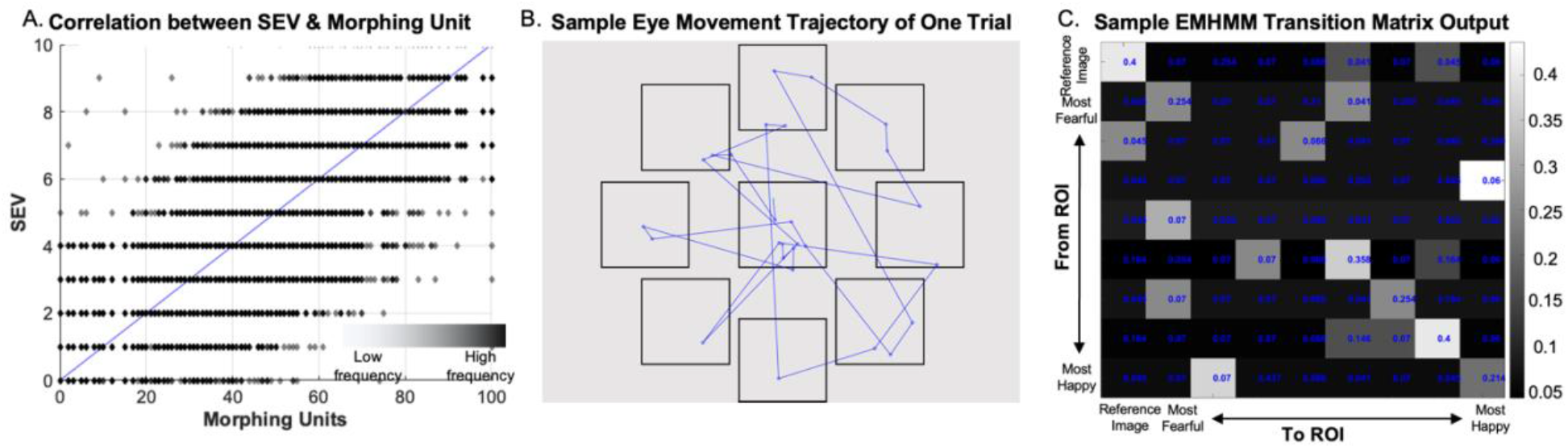
Rating and eye-movement patterns. (**A**) Correlation between SEV and morphing units shows a decent positive correlation. **(B)** Sample participant’s trajectory of eye movement during the 8000-msec in one trial, where the cyan dot indicates the first fixation of that trial. **(C)** Sample EMHMM output of transition matrix, where the transition probabilities have been reordered according to SEV and OEV, so that the top left is the ROI pertaining to the central reference image, next to which is the most fearful ROI of that trial while the bottom right is the happiest ROI of that trial. Each cell indicates the probability of transitioning from a given ROI to another ROI.

In terms of eye measurements, on average, participants exhibited 20.52 fixations (*SD* = .91 fixations) and 4631-msec duration (*SD* =198-msec). **Figure 2B-C** displays a sample participant’s eye movement trajectory in one trial and a sample transition probability matrix generated by EMHMM, respectively.

### Extreme Emotion attracted More Attention

Next, we examined attention allocation as a function of SEV using fixation count, gaze duration, and marginal transition probability. **Figures 3A-C** shows the patterns of the three eye movement measures. There was an ROI Rank Order effect on the percentage of fixation count, *F*(7,224) = 6.93, *p* < .001, η_p_ ^2^ = .18, with a significant quadratic effect, *F*_quadratic_(1,32) = 15.08, *p* < .001, η_p_ ^2^ = .38. Visual inspection revealed a tilted U-shape such that higher proportions of fixation counts were allocated to ROIs with more extreme emotional evidence (outlying) than ROIs with less extreme emotional evidence (inlying). Also, the happiest ROI had the highest percentage of fixation count compared with the inlying ROIs 2 to 7 (*Mean* = 10.50%, all *p* < .05).

**Figure 3.**
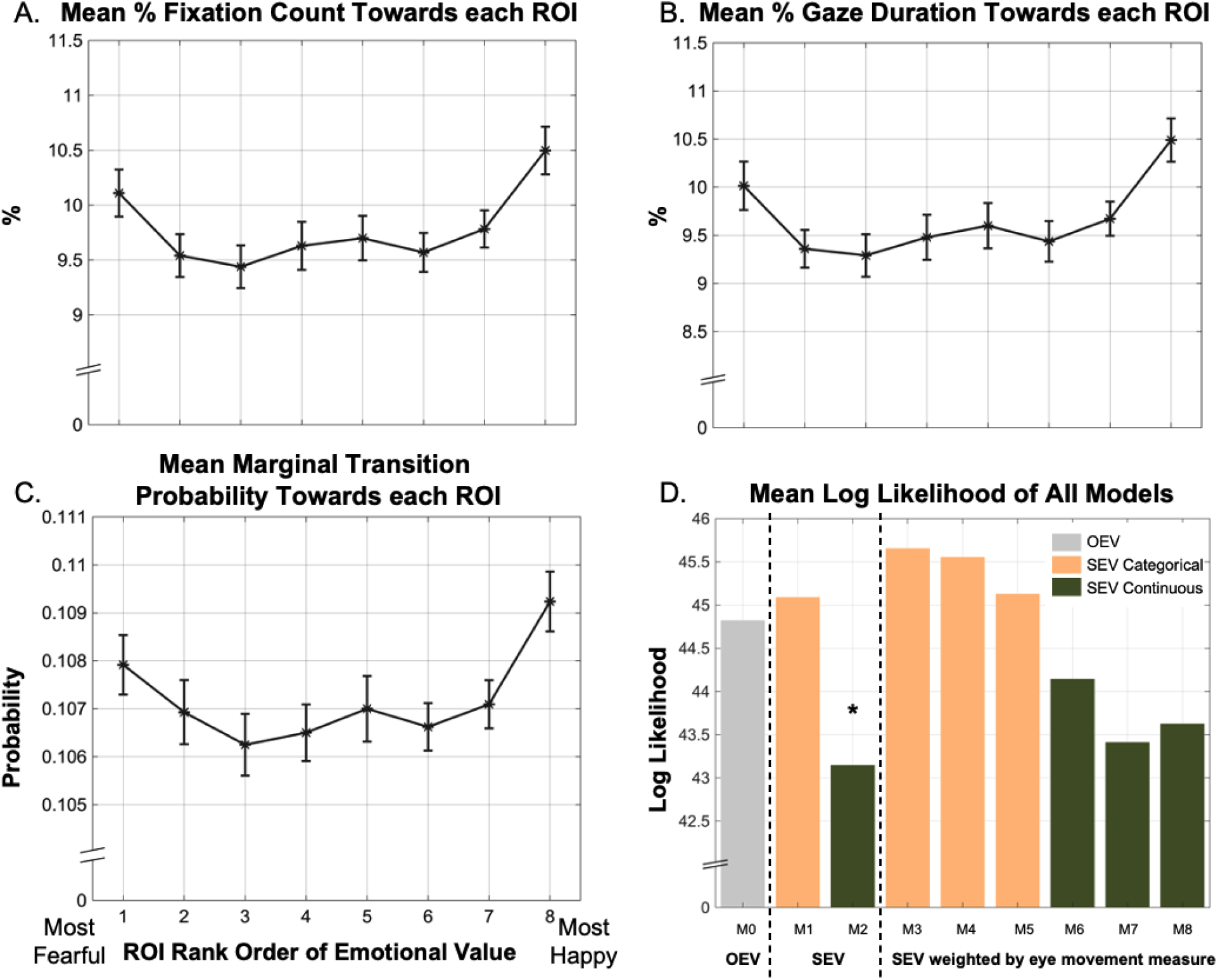
Experiment results. (**A**) The mean fixation count (%), **(B)** gaze duration (%), and **(C)** marginal transition probability towards the eight evidence-bearing face images; the images have been reordered so that 1 indicates the most fearful ROI and 8 indicates the happiest ROI. Error bars represent ±1 SE. **(D)** Mean of each model’s minimized negative log likelihood over all trials. Model 0 uses OEV, whereas Models 1, 3, 4 and 5 use a categorical approach of SEV only (M1), then incorporating Fixation Count (M3), Gaze Duration (M4), Transition Probability (M5) respectively. Models 2, 6 7 and 8 directly use the SEV, taking a continuous approach of SEV only (M2), then weighted with Fixation Count (M6), Gaze Duration (M7), and Transition Probability (M8). The results show that Model 2, where the DV was calculated as a simple average of all SEV had the lowest mean log likelihood (marked by *).

Consistent with the percentage of fixation count, there was an ROI Rank Order effect on the percentage of gaze duration, *F*(7,224) = 7.45, *p* < .001, η_p_ ^2^ = .19, with a significant quadratic effect, *F*_quadratic_(1, 32) = 15.83, *p* < .001, η_p_ ^2^ = .33. Here again, we observed a tilted U-shape in the pattern of gaze duration, supporting that longer gaze durations were allocated to outlying ROIs than inlying ones. The happiest ROI also showed the highest percentage of gaze duration compared with the inlying ROIs 2 to 7 (*Mean* = 10.50%, all *p* < .05).

Finally, marginal transition probability, estimated from the eye movement trajectory of each trial, also revealed an effect of ROI Rank Order, *F*(7,224) = 4.92, *p* < .001, η_p_ ^2^ = .13, with a significant quadratic effect, *F*_quadratic_(1, 32) = 8.82, *p* = .006, η_p_^2^ = .22. On top of the U-shape, the happiest ROI had the highest transition probability compared with the inlying ROIs 2 to 7 (*Mean* = .109, *p* < .05).

We also investigated attention allocation for the first and second halves of the 8000-msec free viewing separately, using fixation count and gaze duration **(Figure 4)**. Percentage of fixation count showed a consistent quadratic effect in both the first epoch, *F*_quadratic_(1,32) = 18.23, *p* < .001, η_p_ ^2^ = .32, and the second epoch, *F*_quadratic_(1,32) = 4.75, *p* = .037, η_p_^2^= .13. The same pattern appeared for gaze duration, showing a consistent quadratic effect in both the first epoch, *F*_quadratic_(1,32) = 19.67, *p* < .001, η_p_ ^2^ = .38 and the second epoch, *F*_quadratic_(1,32) = 5.39, *p* = .027, η_p_ ^2^ = .14.

**Figure 4.**
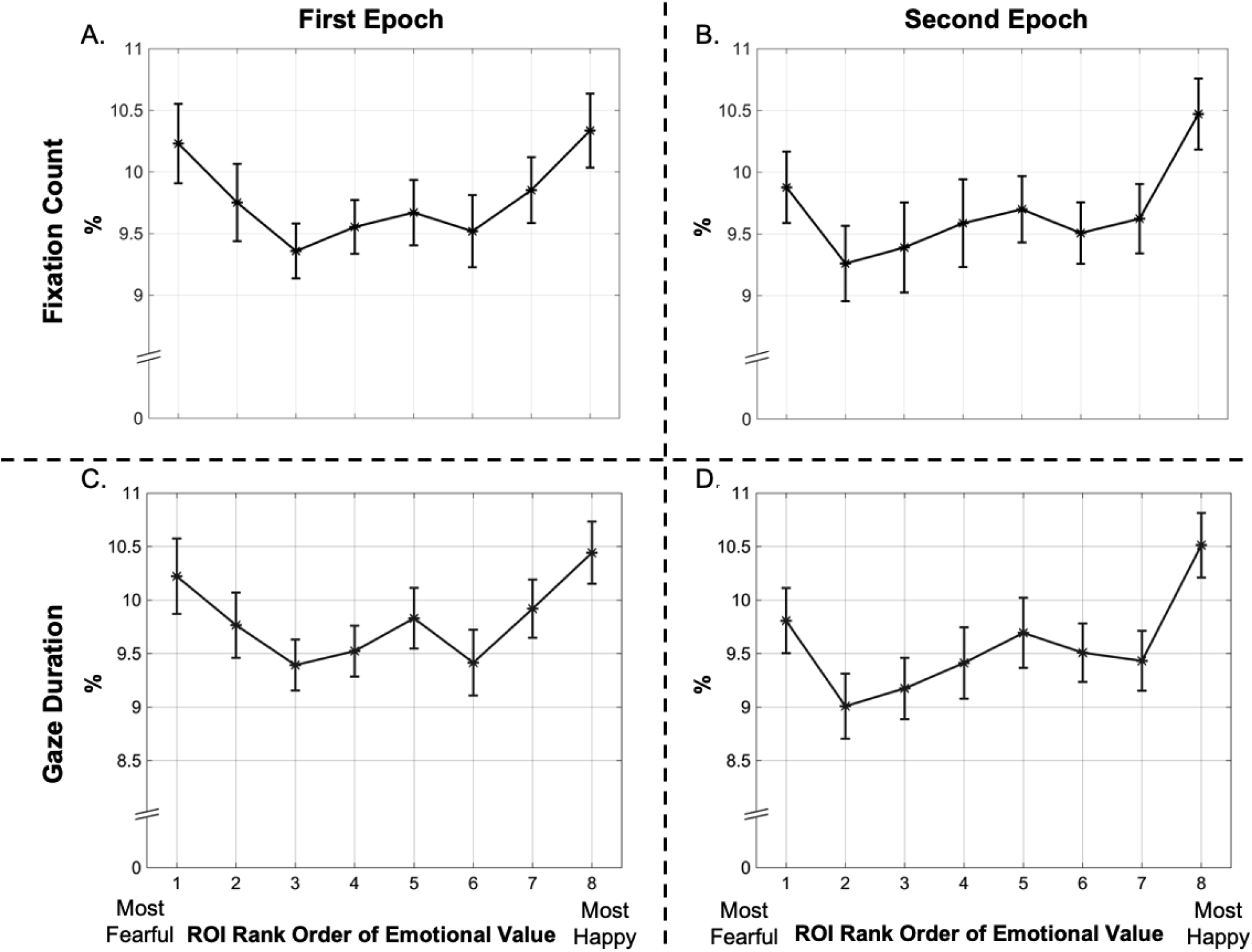
Epoch-based results. (**A**) The mean fixation count (%) to each ROI in the first epoch and **(B)** the second epoch. **(C)** The mean gaze duration (%) in the first epoch and **(D)** the second epoch. The results show a consistent U-shape pattern, where the extreme ROIs were sampled more in both halves of a trial.

Overall, in line with our hypothesis and contrary to indications from non-emotion-related multi-evidence decision-making research, eye movement data demonstrated that extreme emotional evidence attracted more visual attention and were sampled more rather than less. In addition, human participants showed a visual attention bias towards the happiest stimulus.

### Emotional evidence are Weighted Equally

We then examined how pieces of evidence were weighted in evidence integration to form a decision variable. Results **(Figure 3D)** showed that the best fitting-model, the one with the lowest negative log likelihood was M2. In this model, DV was calculated as a simple average across the eight SEVs in each trial. This means that all pieces of evidence were weighted equally. It is noteworthy that models using dichotomized SEV (M1, 3-5) – treating SEV categorically – performed worse than models using SEV as continuous values (M2, 6-8). This shows that decisions are better predicted when the actual magnitude of SEV is taken into consideration. Moreover, the models using a categorical approach also performed worse than the baseline model (M0) which only used OEV and did not take into account any individual differences. This result reiterates the importance of taking the actual magnitude of the weight into consideration. Finally, comparing models with vs without eye movement measures (M1 vs M3-5; M2 vs M6-8) we observe that models weighted by eye movement measures performed worse than the ones using purely SEV. This means that visual attention allocation does not contribute to decisions after considering SEV. Taken together, participants’ decisions were best predicted by weighting all pieces of SEVs equally, with eye movement measures contributing no additional value.

## Discussion

Judging the overall emotional state of a natural scene often necessitates the integration of evidence from multiple sources, a process that requires active sampling of the evidence space and weighting of the sampled emotional evidence. For instance, a speaker assessing whether their audience is enjoying a talk may deliberately scan the expressions of multiple faces and weight such pieces of evidence over several seconds to form a summary impression of the audience’s overall level of engagement or boredom. It is well-known that emotionally salient stimuli draw visual attention, as measured by gaze allocation (Ciesielski et al., 2010; Itti & Koch, 2000; Kollmorgen et al., 2010; Kulke, 2019; Treue, 2003). Most existing studies on emotion perception have presented a single stimulus as the source of evidence (e.g. Mourao-Miranda et al., 2003; Palermo & Rhodes, 2007; Vuilleumier & Pourtois, 2007). For studies using multiple stimuli, evidence is displayed for a very brief duration, which likely engages distributed attention and precludes the examination of how humans intentionally allocate focused attention to sample emotional evidence as well as how such evidence is integrated in the overall decision (e.g. Haberman et al., 2015; Haberman & Whitney, 2010; Ji et al., 2014; Wolfe et al., 2015).

Using eye-tracking and computational modeling, we examined how human participants attentionally sample and weight various pieces of emotional evidence to form a categorical perceptual decision. We examined eye movement data holistically using three types of measures: fixation count, gaze duration and transition probability. Fixation count and gaze duration have been linked to the general importance of a region, as important regions are fixated more than less important ones (Christianson et al., 1991; Jonides et al., 1982; Loftus, 1972; Rayner, 1998). Also, the longer one fixates or gazes at an area, the more information is extracted from it (Caspi et al., 2004; Wu & Kowler, 2013). Lastly, transition probability was estimated using HMM. This approach effectively summarizes the trajectory of visual attention behavior (Chuk et al., 2014). By utilizing these three methods, a comprehensive analysis of the eye movement data was conducted, providing insight into the cognitive and perceptual processes underlying overt attention allocation across stimuli with emotional evidence.

The results revealed that the subjective value of evidence affected how it was sampled, such that more attention was paid to more extreme emotional evidence. This pattern directly contrasts with the general conclusion in decision-making studies utilizing non-emotional stimuli which show less attention to extreme evidence (Bang & Rahnev, 2017; Tsetsos et al., 2012; Vandormael et al., 2017; von Clarenau et al., 2022). The pattern of results in our study may also reflect a key difference in the kind of attention recruited in our task. Given more time to sample the evidence, participants’ attention allocation in our study was guided by their top-down control such that participants were voluntarily allocating their eye gaze across the evidence array. This is different from more rapid previous tasks in which participants may rely on more bottom-up, exogenous capture of attention (Alvarez & Oliva, 2008; Epstein et al., 2020; Haberman & Whitney, 2010). In addition, we found evidence for positive bias compared to negative bias during sampling, which is consistent with the general attention literature (Pool et al., 2016). Overall, our eye-tracking findings highlight the distinction in evidence sampling in emotion versus non-emotion related decision-making studies indicating that humans used a biased sampling strategy by purposefully sampling emotional evidence that is more extreme.

In terms of evidence weighting, previous research using two-alterative tasks has shown that the attended stimulus is weighted higher in decisions (Krajbich et al., 2010; Orquin et al., 2021; Thomas et al., 2019; Vandormael et al., 2017). Hence, we hypothesized that biased sampling (towards the most extreme emotional evidence) will translate into a biased weighting in the overall decision. In contrast to this hypothesis, we found that while more attention was allocated to extreme emotional evidence, higher attention did not translate into greater weighting in the overall decision. In fact, the best-fitting model was the simple mean subjective value model, indicating that participants weighted all pieces of emotional evidence equally. This is in line with previous research showing that humans are sensitive to the mean emotion even with a glance (Haberman et al., 2015; Haberman & Whitney, 2010; Whitney & Yamanashi Leib, 2018). Then, given that there was no feedback or reinforcer associated with either decision, the optimal decision-making strategy in our study was indeed to simply average across all pieces of emotional evidence, an effect that has been seen in earlier studies with non-emotional stimuli too (Brown & Heathcote, 2005; Smith & Ratcliff, 2004; Van Den Berg & Ma, 2012).

While attentional biases may be adaptive by allowing us to sample evidence of threat and reward in our environment, our overall decision making is not likely to be adaptive if it is heavily weighted in favor of extreme evidence that captures our attention. A decision-making strategy that avoids overweighting extreme evidence will facilitate individuals to interact with the environment in an adaptive manner by forming accurate judgments. Here, we show that humans tend to sample the evidence space biased by emotional saliency, especially positivity, but give equal weights across all pieces of evidence. However, it remains to be examined if too much attention allocated to extreme emotional evidence translates into biased weighting in clinical conditions like anxiety or depression. For instance, anxiety is characterized by hypervigilance (Grupe & Nitschke, 2013; Power & Dalgleish, 2015), which may manifest in an increase in threat-oriented attention biasing the perceptual evidence sampling and an overweighting of threatening information. Future research examining whether individuals with high anxiety may allocate more attention and even more weight to negative evidence can facilitate our understanding of the related psychopathology.

The dynamics between several brain regions involved in attention, value-based decision-making, and saliency processing, may play critical roles in the current task. First, the encoding of subjectively valuated emotional evidence may involve the amygdala, the ventromedial prefrontal cortex (vmPFC) and ventral striatum, as these regions are known for processing salient and subjectively valuated information (Anderson & Phelps, 2001; Blair, 2008; Jin et al., 2015; LeDoux, 1998; Levy & Glimcher, 2012; Moratti et al., 2004; Padoa-Schioppa & Assad, 2006; Zhang et al., 2017). Active sampling of emotional evidence through eye movement may involve the lateral intraparietal area, which is known to be a hub for directing overt visual attention and integrating visual input driven by values in decision making (Louie et al., 2011; Mohanty et al., 2009; Reynolds & Desimone, 2003). Dorsolateral prefrontal cortex (dlPFC) is likely involved in guiding the purposeful sampling (Gray et al., 2002). Similarly, the cingulate cortex is proposed to play a key role in integrating emotion and cognition, thus it may be associated with guiding attention during evidence sampling based on emotional value (Mohanty et al., 2007; Shackman et al., 2011). Finally, the output of accumulated evidence may be passed down to anterior part of the prefrontal cortex (PFC) for evidence weighting (i.e., decision-value formation) and generating binary decisions (Gold & Shadlen, 2007; Hanks & Summerfield, 2017). Taken together, the attention network, regions encoding saliency as well as valuation may play roles in sampling and integrating emotional evidence based on subjective valuation.

This study has certain limitations. While this study aimed to cater toward the naturalistic setting of emotion-related perceptual decision making, the multiple stimuli used within each trial were of the same face actor/actress, which is unrealistic. This design was chosen to ensure that the experiment was highly controlled and to address the dynamic property of facial expressions within an individual. Our findings provide foundations for further exploration such as deliberate decision making regarding a crowd of faces from multiple identities (Elias et al., 2017). Also, the current study focused on “happy” as the “positive” valence emotion, and “fearful” as the “negative” valence emotion, because they are of opposite valence but similar arousal level. Future research may examine whether the current findings can be generalized to different categories of anchoring emotions as well as more ambiguous emotions (e.g., surprise). Moreover, the current study recruited predominantly Asian participants and used Asian face actors as stimuli as this is indeed the local sociocultural context of the institution where the experiments were held. Future research can explore these experiments in other countries and demographics.

## Concluding remarks

Decision making regarding emotional information has been a long-standing interest in affective and cognitive sciences. Findings from the present study elucidate how humans integrate multiple pieces of motivationally inherently salient evidence to form an overall perceptual judgment and highlight the distinctive process adopted in deliberate decision-making regarding emotional stimuli. Also, the psychological and computational mechanisms uncovered in the present study shed light on how emotion-related decision making occurs generally and provide clues into how it may be biased in people with emotion-related psychopathology.

## Acknowledgement

The authors thank Dr. Antoni B Chan and Anita Liao for their help in eye movement data management and thank the research assistants who have helped data collection.

## Preregistration and open science

We pre-registered the study on the Open Science Framework (OSF). Pre-registrations, materials are available at https://osf.io/e8jvb. Data can be made available upon request to the corresponding author.

